# Generation of a SARS-CoV-2 reverse genetics system and novel human lung cell lines that exhibit high virus-induced cytopathology

**DOI:** 10.1101/2023.03.08.531833

**Authors:** Juveriya Qamar Khan, Megha Rohamare, Karthic Rajamanickam, Kalpana K Bhanumathy, Jocelyne Lew, Anil Kumar, Darryl Falzarano, Franco J Vizeacoumar, Joyce A Wilson

## Abstract

The global COVID-19 pandemic continues with an increasing number of cases worldwide and the emergence of new SARS-CoV-2 variants. In our study, we have developed novel tools with applications for screening antivirals, identifying virus-host dependencies, and characterizing viral variants. Using reverse genetics, we rescued SARS-CoV-2 Wuhan1 (D614G variant) wild type (WTFL) and reporter virus (NLucFL) using molecular BAC clones. The replication kinetics, plaque morphology and titers were comparable between rescued molecular clones and a clinical isolate (VIDO-01 strain), thus providing confidence that the rescued viruses can be used as effective replication tools. Furthermore, the reporter SARS-CoV-2 NLucFL virus exhibited robust luciferase values over the time course of infection and was used to develop a rapid antiviral assay using remdesivir as proof-of-principle. In addition, as a tool to study lung-relevant virus-host interactions, we established novel human lung cell lines that support SARS-CoV-2 infection with high virus-induced cytopathology. Six lung cell lines (NCI-H23, A549, NCI-H1703, NCI-H520, NCI-H226, and HCC827) and HEK293T cells, were transduced to stably express ACE2 and tested for their ability to support virus infection. A549^ACE2^ B1 and HEK293T^ACE2^ A2 cell lines exhibited more than 70% virus-induced cell death and a novel lung cell line NCI-H23^ACE2^ A3 showed about ∼99% cell death post-infection. These cell lines are ideal for assays relying on live-dead selection and are currently being used in CRISPR knockout and activation screens in our lab.

**Importance:** We used a reverse genetics system to generate a wild type as well as a nanoluciferase-expressing reporter clone of SARS-CoV-2. The reporter virus allows for rapid transient replication assays and high throughput screens by detection of virus replication using luciferase assays. In addition, the reverse genetic system can be used to generate mutant viruses to study phenotypes of variant mutations. Additionally, unique human lung cell lines supporting SARS-CoV-2 replication will aid in studying the virus in a lung-relevant environment and based on high cytopathology induced in some cell lines, will be useful for screens that rely on virus-induced cell death for selection. Our study aims to enhance and contribute to the current replication tools available to study SARS-CoV-2 by providing rapid methods, virus clones and novel lung cell lines.

## Introduction

Severe acute respiratory syndrome coronavirus 2 (SARS-CoV-2) is responsible for the global pandemic of COVID-19 disease and has spread around the world rapidly since its discovery in 2019. It was first reported in Wuhan, the capital city of Hubei province in China and since then has led to over 6 million deaths globally (as of January 2023) [WHO Coronavirus (COVID-19 Dashboard) (WHO, 2020)]. The virus affects the respiratory tract, and the most characteristic symptoms of COVID-19 are flu-like symptoms, such as fever, fatigue, dry cough, and loss of taste or smell. In severe life-threatening cases, patients with inflammation of the lungs can experience trouble breathing, persistent pressure in the chest, confusion and require life support (1). The primary transmission mode is direct contact with the infected individual or by sneezing and coughing, leading to the transfer of nasal droplets (2). There have also been cases of asymptomatic COVID-19 which led to the silent transmission of the disease (3).

SARS-CoV-2 is a positive sense single-stranded RNA virus belonging to the genus *Betacoronaviridae*. The genome of SARS-CoV-2 is ≈ 30 kb in length and encodes structural, non-structural, and accessory proteins (4). The genome has a 5’ cap and a 3’ poly-A sequence and the 5’ and 3’ untranslated regions (UTR) of the genome are highly structured. Two-thirds of the genome encodes a single polypeptide called ORF1ab and consists of non-structural proteins. ORF1a encodes polypeptide 1a (pp1a), similarly, ORF1b encodes polypeptide 1b (pp1b). Polypeptide 1ab is a result of ribosomal frameshift during the translation of ORF1a. These polyproteins are processed by viral proteases and give rise to 16 non-structural proteins involved in the replication and transcription (5). SARS-CoV-2 has four structural proteins-Spike (S), Membrane (M), Envelope (E) and Nucleocapsid (N). The function of the S protein is to bind to the host cell surface receptor, angiotensin-converting enzyme 2 (ACE2) and mediate virus entry. ACE2 is an integral membrane protein and serves as a receptor for SARS-CoV-2 by binding to the receptor binding domain of S protein (6). The virus shares more than 80% nucleotide similarity with the SARS-CoV genome and more than 90% similarity in essential structural elements and proteins (5). The evolution of the SARS-CoV-2 genome has given rise to various variants, with several variants of concern (VOC) sequentially becoming dominant worldwide over the course of the pandemic [WHO Coronavirus (COVID-19) Dashboard, (WHO, 2020)]. While rapid development and distribution of SARS-CoV-2 vaccines have reduced the risk of hospitalization, few treatments exist for those with severe disease. Monoclonal antibody therapies were effective in treating early SARS-CoV-2 variants but most of these are no longer effective against Omicron due to evolution of S mutations and immune escape. Small molecule drug therapies are also currently limited to remdesivir (Veklury®), a combination of nirmatrelvir and ritonavir (brand name Pfizer-Paxlovid^™^), and molnupiravir (Lagevri^™^) but the supply of these treatments is limited, and drug-drug interactions preclude their widespread use [Government of Canada, (Canada, 2022)]. Cases of COVID-19 are still on the rise due to waning antibody response post-vaccination, and immune escape of emerging variants. Effective antivirals for SARS-CoV-2 can be developed based on a better understanding of virus replication and the development of novel virus and cell culture systems, including viruses with easy-to-monitor reporters. In this study, we have designed and constructed wild-type and modified genomes using reverse genetics and used them to generate wild-type and reporter viruses and a subgenomic replicon. We have also generated novel human cultured lung cells that are highly susceptible to SARS-CoV-2. These cell lines exhibited high cytopathic effects which are ideal for assays that use live-dead cell selection such as CRISPR knockout screens to identify host dependency factors. We also show the usefulness of the reporter virus and cells in antiviral drug screens. Together these tools provide models for research to understand the mechanism of viral infection, discover new antiviral strategies and screen effective antivirals.

## Results

### Design and recovery of infectious viruses derived from SARS-CoV-2 molecular clones

A SARS-CoV-2 wild type full-length (WTFL) synthetic genome based on the ancestral Wuhan1 virus sequence (Wuhan-1 NC_045512) and including the Spike D614G mutation was ordered from Codex DNA Inc. in a BAC vector. The clone included a T7 RNA polymerase promoter sequence immediately upstream of the 5’ end of the genome and a unique SbfI restriction site downstream of an encoded poly (A) tail at the 3’ end (Figure 1Ai). We also designed and ordered a NLuc reporter version of this clone, SARS-CoV-2-NLucFL in which a NLuc gene replaced ORF7a (7). This clone was used to generate an infectious reporter virus for rapid analysis of SARS-CoV-2 replication (Figure 1Aii). The sequences were synthesized and assembled by Codex DNA, Inc., who provided us with purified bacmid DNA based on our sequence design. To use the plasmids to recover SARS-CoV-2 WTFL and SARS-CoV-2 NLucFL viruses, genomic RNA was transcribed *in vitro* and co-electroporated into Vero76 cells with N mRNA (7). By 3 days post-electroporation the cells developed significant cytopathic effects (CPE), observed as cell rounding, and cell death similar to the CPE seen with the SARS-CoV-2 Wuhan1 clinical isolate infection (Figure 1B). The supernatant from this stock was stored as passage 0 (P0) and was further amplified in Vero76 cells to make P1 stock which was used as working stock for our experiments.

**Figure 1:**
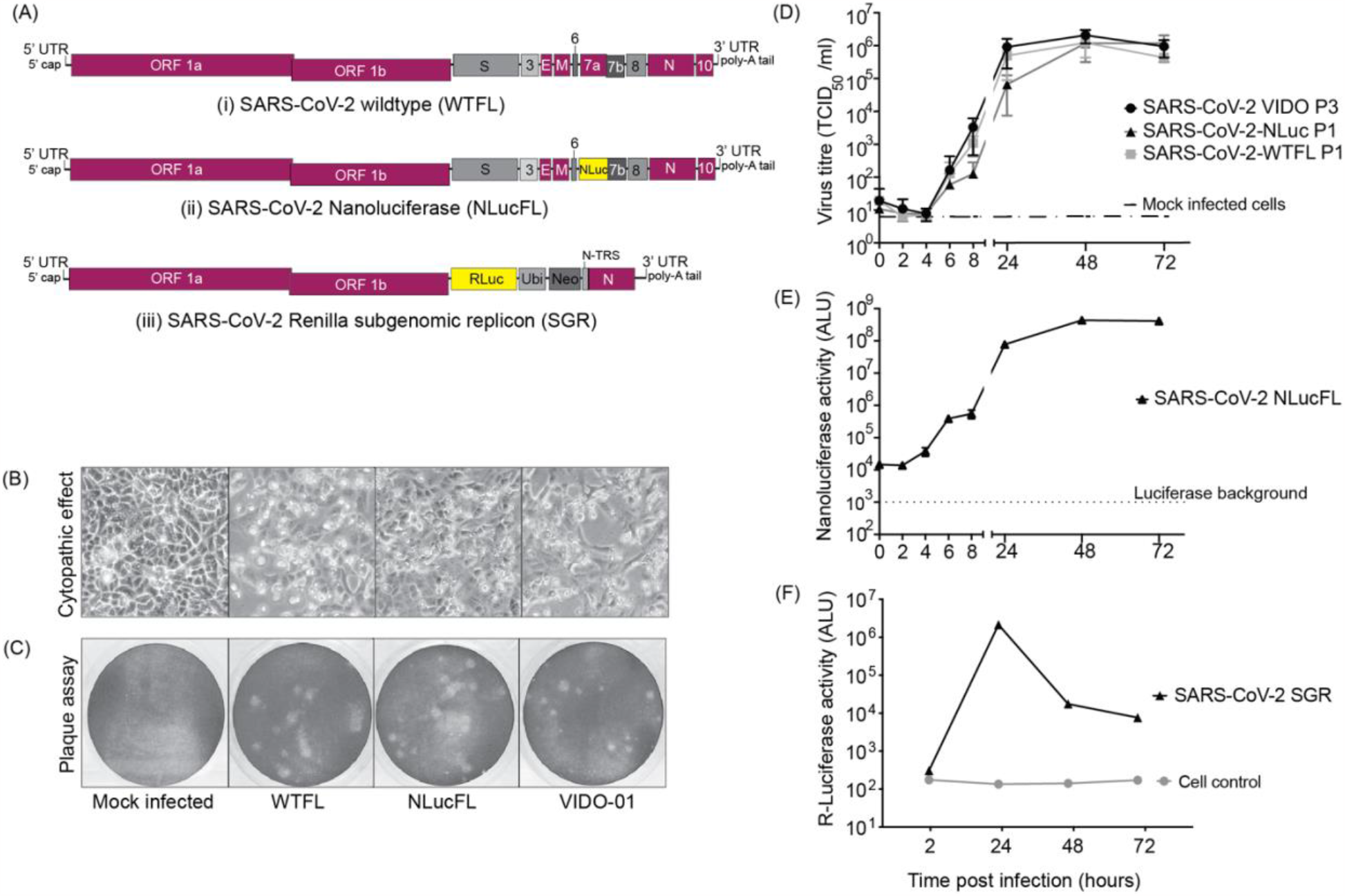
SARS-CoV-2 molecular clones and rescued viruses. (A) The synthetic molecular clones for wild type full-length virus (WTFL) (i), NLuc expressing reporter virus (NLucFL) (ii), and sub-genomic replicon (SGR) (iii) are represented schematically. In the clone BAC plasmid, the genome is preceded by a T7 polymerase and a poly A sequence at the 3’ end. In NLucFL, the NLuc gene replaces ORF7a and in SGR, Renilla luciferase (RLuc)-Ubiquitin (Ubi)-neomycin resistance gene (Neo) replace genes S-to ORF8 and are under control of the S-TRS. The various genes of the viral genome are labelled as; L-leader sequence, ORF-open reading frames, S-spike, E-envelope, M-membrane, N-nucleocapsid, and UTR-untranslated region. (B) Virus-induced CPE (defined by cell rounding and detachment from monolayer or cell death) and (C) plaques observed in Vero76 cells 2-3 days post-infection were comparable in all the clones and the clinical isolate wild type VIDO-01 virus. (D) Replication kinetics of rescued SARS-CoV-2 WTFL and NLucFL, were compared to SARS-CoV-2 wild-type lab-isolated strain VIDO-01. The data is an average of three independent experiments and error bars represent the standard deviation. ALU: arbitrary luminescence units. (E) NLuc expression kinetics were observed to be robust, with peak values observed on Day 2 post-infection. The data is an average of three independent experiments and error bars represent the standard deviation. (F) Transient replication assays with SARS-CoV-2 SGR revealed peak *Renilla* luciferase expression on Day 2 post-electroporation in Huh7.5 cells. Data are representative of three independent experiments.

### Characterization of infectious virus recovered using reverse genetics

To characterize the virus obtained from molecular clones in comparison to the clinical VIDO-01 isolate, we performed plaque assays in Vero76 cells with SARS-CoV-2 VIDO-01 (P3), WTFL (P1) and NLucFL (P1) virus stocks. The morphology of plaques generated by the clone-derived SARS-CoV-2 WTFL and -NLucFL viruses were slightly larger (Figure 1C) and uniform when compared to the plaques made by the SARS-CoV-2 VIDO-01 virus which were of varying sizes (Figure 1C). This was expected since the recovered virus is from a single molecular clone whereas the clinical isolate may contain several quasi-species. The virus titres obtained from VIDO-01 P3, WTFL P1 and NLucFL P1 were comparable in the range of 1-3 × 10^6^ TCID_50_units/mL.

To evaluate the efficiency of the reverse genetics system and assess the competency of the recovered infectious viruses as successful replication tools, we compared the replication kinetics over a period of three days between the infectious clones and the VIDO-01 clinical isolate (Figure 1D). The SARS-CoV-2-WTFL, -NLucFL and -VIDO-01 clinical isolates showed comparable growth trends, with the virus titers reaching a plateau, and slightly decreasing post-48 hours (Figure 1D). This suggests that the cloned viruses and the insertion of the NLuc gene did not significantly affect replication kinetics or virus titers. Since the reporter virus NLucFL showed similar replication kinetics (Figure 1D), we further evaluated the reporter expression of SARS-CoV-2 NLucFL over a period of 72 hours. The NLuc expression showed a similar trend compared to the virus titers reaching a peak at 48 hours (Figure 1E). The exponential luciferase expression kinetics and similarity in growth curves generated based on virus titres and NLuc expression support that NLuc expression can be used as a surrogate measure of virus replication in virus replication assays.

### Design and assessment of transient replication of SARS-CoV-2 non-infectious sub-genomic replicons

To develop a SARS-CoV-2 replication system that can be used in a BSL2 laboratory, we generated a non-infectious sub-genomic replicon (SGR) clone by deleting all the structural and accessory genes in the virus genome, except for the N gene which is required for virus replication. In place of the structural genes and upstream of the N sequence, we added a Renilla luciferase gene to assess replication of the SGR, and a neomycin resistance gene to enable the selection of stable SARS-CoV-2 expressing cell lines, separated by a ubiquitin sequence. (Figure 1Aiii). This cassette was under the regulation of the transcriptional regulatory sequence of S protein (S-TRS). The purified DNA obtained from Codex DNA, Inc. was used to prepare *in vitro* transcribed SGR RNA that was co-electroporated in cells with an equal concentration of N mRNA. We were successful in measuring transient RLuc expression twice in Huh 7.5 cells and observed that the luciferase expression peaked at 24 hours post-electroporation (Figure 1F). We also observed CPE starting at 48 hours post-electroporation and a sharp decrease in luciferase expression at the same time. Attempts to detect transient SGR replication in other lung cell lines such as NCI-H226 and A549 were not successful, and we were also not able to isolate stable SARS-CoV-2 expressing cells following cell selection with G418 In general, we found transient replicon replication assays to be difficult to perform and success rates were not reliable enough to warrant its use for routine replication assays in BSL2.

### Establishing rapid tools for antiviral assays using remdesivir

Based on robust replication and ease of use of the SARS-CoV-2 NLucFL reporter virus we tested and optimized its use in high-throughput replication and inhibitor screening assays. To test the reproducibility of antiviral inhibition of our molecular clones in comparison to SARS-CoV-2 VIDO-01 clinical isolate, we first tested the antiviral effect of remdesivir on WTFL, NLucFL and VIDO-01 viruses in Vero76 cells based on a dose-dependent reduction in virus titers. The cells were infected at 0.01 MOI for 1 hour, followed by compound treatment at various concentrations for 48 hours. A dose-dependent inhibition curve was generated by calculating virus titers using TCID_50_ (Figure 2A-C). The resulting 50% Effective concentration (EC_50_) of remdesivir for VIDO-01, WTFL and NLucFL were comparable to each other at 4.95, 4.65 and 6.55 µM respectively, providing confidence in using the molecular clones as effective replication tools (Figure 2A-C). Furthermore, to optimize the reporter assay and test NLucFL as a rapid assessment proxy system for drug screening, NLuc expression was assessed at various remdesivir concentrations in parallel. NLuc expression was also observed to decrease in a dose-dependent manner giving an EC_50_of 6.15 µM (Figure 2D). The CC_50_for remdesivir in Vero76 cells in our assay was 183.7 µM resulting in the selective index (SI) (CC_50_/EC_50_) to be 37.11, 39.5, 28.04 for SARS-CoV-2 VIDO-01, -WTFL and -NLucFL virus, respectively. The SI for NLucFL based on the luciferase assay was 29.86. Thus, the NLuc reporter virus system can be used in high-throughput drug screens.

**Figure 2:**
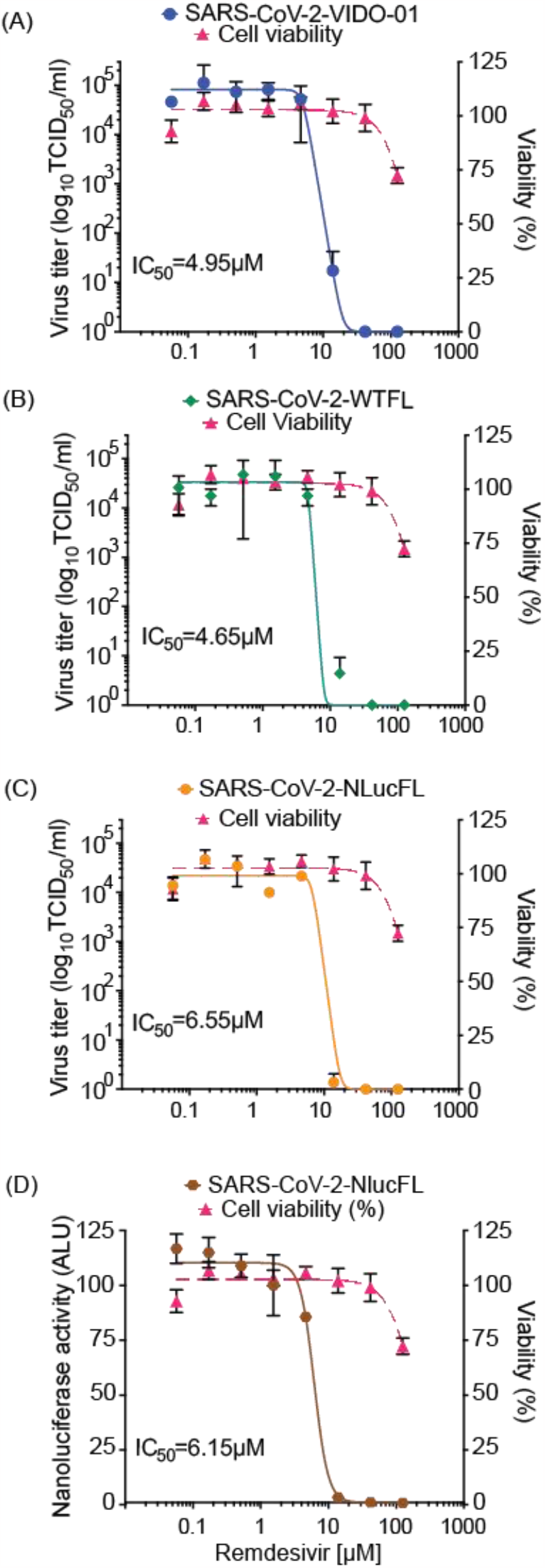
Antiviral inhibitor assays using SARS-CoV-2 WTFL and NLuc reporter viruses derived from the molecular clones. Dose-dependent effect of remdesivir was assessed on replication of (A) SARS-CoV-2 wild type VIDO-01 isolate (EC_50_= 4.95µM), (B) SARS-CoV-2 WTFL (EC_50_= 4.65µM), (C) SARS-CoV-2 NLucFL (EC_50_= 6.55µM), using virus titrations and with (D) SARS-CoV-2 NLucFL (EC_50_= 6.15µM) using the NLuc reporter expression as a surrogate for virus replication. The CC_50_was calculated to be 183.7µM. Cell viability was assessed in each assay and % viability was read, depicted on the right y-axis. The data is an average of three independent experiments and error bars represent the standard deviation.

### Selecting highly susceptible lung cell lines showing virus-induced cytopathic effect

To study SARS-CoV-2 replication and virus-host interactions convenient cell culture systems are required, and to study tropism for the lungs a lung cell culture system is desirable. In addition, many genome-wide screening systems such as CRISPR knockout or activation screens rely on virus-induced cell death for the selection of cells susceptible or not to SARS-CoV-2 infection. SARS-CoV-2 infects Calu3 cells, but this cell line is inconvenient for many types of analyses due to their slow growth and is a poor choice for genome-wide screens because of low levels of CPE (8). Our goal was to develop lung cell lines that are susceptible to SARS-CoV-2 infection. To this end, we screened eight cell lines, including A549, HEK293T and five novel lung cells-NCI-H23, NCI-H1703, HCC827, NCI-H520 and NCI-H226 to test if they support virus infection, and found that none of them exhibited SARS-CoV-2 infection-induced CPE. For successful infection in the host, SARS-CoV-2 requires the cell surface receptor ACE2, which also dictates virus tropism. To increase virus susceptibility, we transduced the cell lines with human ACE2 lentiviruses and generated cell lines that stably expressed ACE2, and re-evaluated CPE caused by SARS-CoV-2 VIDO-01 infection. Virus-induced CPE was analyzed based on cell viability for 3 days post-infection using Promega Viral ToxGlo^™^ assay (Figure 3-4, panel I) and observed visually by microscopy (Figure 3-4, panel III). Cell surface ACE2 expression was also assessed by flow cytometry (Figure 3-4, panel II).

**Figure 3:**
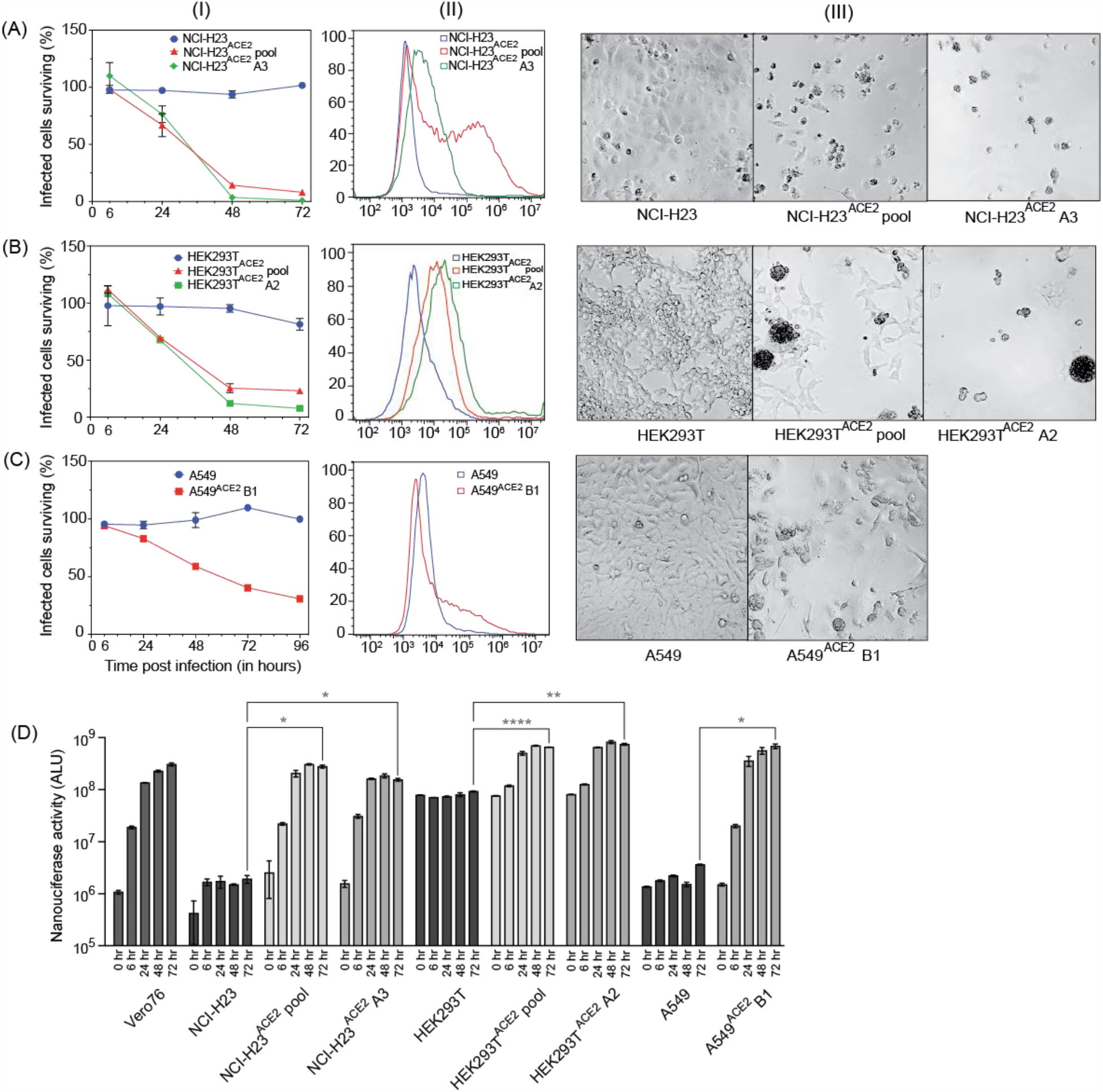
Three cell lines successfully transduced with ACE2, supporting virus infection, and showing high virus-induced CPE. For figure 3A, -B, -C, Panel (I) depicts cell viability quantified post-infection with SARS-CoV-2 VIDO indicating virus-induced CPE; Panel (II) depicts flow cytometry analysis to assess the expression of ACE2; Panel (III) depicts a visual representation of CPE post-infection with SARS-CoV-2 VIDO-01 as seen under the microscope, for the following cell lines (A) NCI-H23, NCI-H23^ACE2^ cell pool, and NCI-H23^ACE2^ clone A3; (B) HEK293T, HEK293T^ACE2^, and HEK293T^ACE2^ clone A2; (C) A549, and A549^ACE2^ clone B1. (D) Replication kinetics of SARS-CoV-2 NLucFL virus in untransduced and ACE2 transduced cell lines assessed by expression of NLuc. The data is an average of three independent experiments and error bars represent the standard deviation. Statistical significance was determined on the 72-hour values using one-way ANOVA; *P <0.0332, **P <0.0021, ***P <0.0002, ****P <0.0001.

### NCI-H23^ACE2^ monoclonal cell line showed the highest virus susceptibility at 99% virus-induced cell killing

One of our goals was to develop a cell line that will be useful for screens, such as CRISPR screens, that rely on live-dead selection to analyze SARS-CoV-2 virus-host interactions. To that end, the best cell line was NCI-H23^ACE2^ which showed only 15% surviving cells 72 hours post-infection (Figure 3A-I). Cell clones were generated from this cell pool and 5 clones were screened for infection and CPE. NCI-H23^ACE2^ clone A3 showed only 1% surviving cells post SARS-CoV-2 infections (Figure 3A-I, III). Flow cytometry shows a sharp peak of ACE2 expression in the monoclonal cell line vs a broader peak in the transduced cell pool (Figure 3A-II) and suggests the high CPE is likely due to efficient receptor expression and virus infection of the cells. Thus, NCI-H23^ACE2^ cells are highly susceptible to SARS-CoV-2 infection, show a remarkable 99% virus-induced CPE, and are thus ideal for screens that rely on virus-induced cell death. These screens are currently underway.

### Additional monoclonal cell lines identified to support robust virus replication

Similarly, ACE2 transduced HEK293T, and A549 cells also supported SARS-CoV-2 infection and exhibited high virus-induced cell death. SARS-CoV-2-VIDO-01 infection of ACE2 transduced HEK293T^ACE2^ cells resulted in only 23% and 10% surviving cells 72 hours post-infection in cell pools and monoclonal cell lines (clone A2) respectively (Figure 3B-I). ACE2 expression was assessed by flow cytometry and correlated with susceptibility to infection (Figure 3B-II), and SARS-CoV-2-induced CPE in these cells was defined by cell clumping, rounding and cell death (Figure 3B-III).

Infection of monoclonal A549^ACE2^ cells (clone B1) resulted in 68% virus-induced cell death (31% surviving cells) 4 days (96 hours) post-infection by the SARS-CoV-2 VIDO-01 isolate (Figure 3C-I) and CPE was defined by syncytia formation, cell rounding and death (Figure 3C-III). Although the ACE2 peak shift in flow cytometry was less pronounced in the A549^ACE2^ transduced cells (Figure 3C-II), an extended tail on the x-axis for the ACE2 transduced monoclonal cell line can be observed indicating positive ACE2 expression.

Additionally, we used the NLucFL reporter virus to study the replication kinetics over a period of 3 days post-infection of the untransduced cells and their transduced counterparts (Figure 3D). Vero76 cells, which support robust virus replication, were used as a positive control to assess a typical growth curve. The ACE2 untransduced cell lines exhibited basal levels of NLuc expression as can be seen with NCI-H23, HEK293T and A549 cells (Figure 3D), and the ACE2 transduced cell pools and monoclonal cell lines exhibited an increase in the NLuc expression from 0-72 hours post-infection (Figure 3D), thus confirming that the cell lines successfully support virus replication. NCI-H23^ACE2^ cells have also been used successfully for antiviral drug testing against SARS-CoV-2 (9).

### Screening of other lung cell lines with less pronounced virus-induced CPE

A secondary aim was to develop cell lines that support SARS-CoV-2 infection but display lower levels of CPE and three ACE2 transduced human lung cell lines showed this phenotype. NCI-H1703^ACE2^ transduced cells showed 82% virus-induced CPE, defined by cell rounding and cell death (Figure 4A-I). Flow cytometry for these cell lines also confirmed ACE2 expression (Figure 4A-II). HCC827^ACE2^ and NCI-H520^ACE2^ exhibited less than 50% cell death at most even at 4 days post-infection, (Figure 4B, C-I) and 30% CPE was seen for NCI-H226^ACE2^ (Figure 4D-I) at day 6 post-infection. The ACE2 expression assessed by flow cytometry can be seen in figure 4-panel II and cell death was observed by microscopy as seen in figure 4-panel III. Although unable to kill most cells by SARS-CoV-2 VIDO-01 infection, the ACE2 transduced cells show a moderate increase in luciferase expression over a period of 3 days after infection with the SARS-CoV-2 NLucFL virus (Figure 4E). The untransduced cells showed basal levels of luciferase throughout the course of 3 days (Figure 4E). Interestingly, HCC827 showed increasing trends in luciferase expression indicating virus replication but failed to show visible signs of CPE (Figure 4E, B).

**Figure 4:**
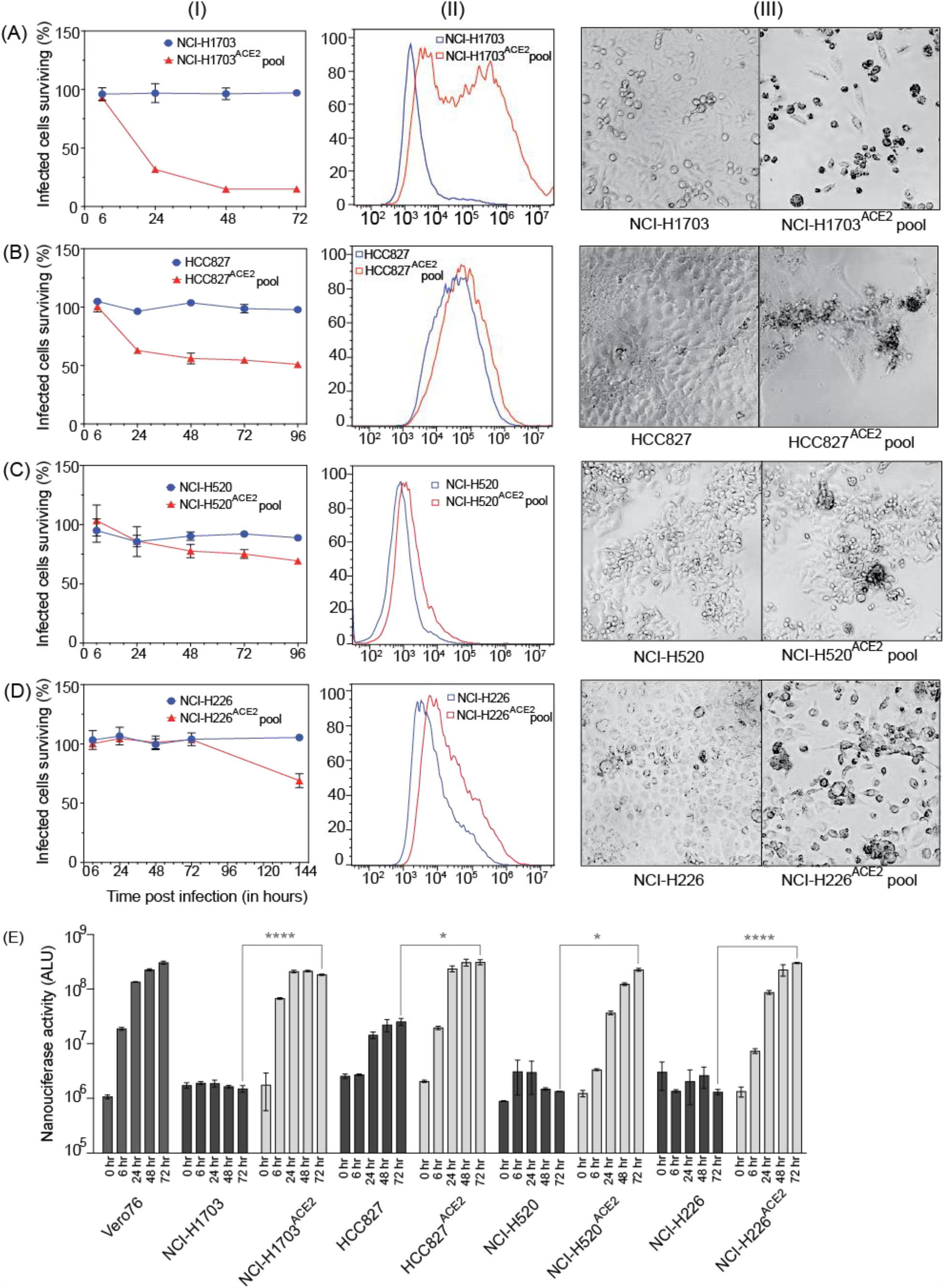
Lung cell lines transduced with ACE2, supporting virus infection at varying levels. Cell lines (A) NCI-H1703, and NCI-H1703^ACE2^ cell pool; (B) HCC827, and HCC827^ACE2^ cell pool; (C) NCI-H520 and NCI-H520^ACE2^ cell pool; (D) NCI-H226, and NCI-H226^ACE2^ cell pool were analyzed for their ability to support SARS-CoV-2 infection with and without ACE2 transduction. Panel (I) quantifies cell viability after infection with SARS-CoV-2 VIDO-01. Panel (II) depicts flow cytometry analysis to assess the expression of ACE2. Panel (III) is a light micrograph of the cells infected with SARS-CoV-2 VIDO-01 to show CPE. (D) shows replication kinetics of SARS-CoV-2 NLucFL virus in untransduced and ACE2 transduced cell lines assessed by expression of NLuc. The data is an average of three independent experiments and error bars represent the standard deviation.. Statistical significance was determined on the 72-hour values using one-way ANOVA; *P <0.0332, **P <0.0021, ***P <0.0002, ****P <0.0001.

In conclusion, we have developed a range of virus clones and ACE2 expressing human lung cell lines to be shared and used for various applications to study SARS-CoV-2 and accelerate the development of antiviral therapeutics.

## Discussion

In this study, we report the generation of virus clones and cell lines as replication tools to study SARS-CoV-2 virus-host interactions and rapidly screen for antiviral agents. We have successfully used reverse genetics to generate wild type and reporter SARS-CoV-2 viruses. Using synthetic cDNA clones of SARS-CoV-2 sequences followed by *in vitro* RNA transcription, we recovered wild type and recombinant viruses in Vero76 cells and successfully characterized them. We compared the wild type full-length clone (WTFL) and the Nluc expressing reporter clone (NlucFL) with the SARS-CoV-2 VIDO-01 clinical isolate and the resulting CPE, plaque morphology and replication kinetics between the three viruses were found to be comparable (Figure 1B-D). This highlights that the virus obtained from molecular clones recapitulates the replication properties of the wild type clinical isolate and can be used as equivalent to clinical isolates for *in vitro* studies on SARS-CoV-2. Our system differs from others in that it required us to simply order fully assembled SARS-CoV-2 genomes which were used to generate viruses without having to amplify the DNA in bacteria or assemble multiple fragments as is required for other SARS-CoV-2 reverse genetic systems (7, 10, 11). This strategy is beneficial during pandemics because it allows virology labs to generate wild-type, mutant, or reporter viruses quickly using only sequence information.

We have also confirmed the potential to use the reporter virus to study virus replication kinetics and drug screens. NLuc expression during replication of the reporter virus SARS-CoV-2 NLucFL corresponded to its replication kinetics indicating that the luciferase assay can be used as a proxy system to detect virus replication (Figure 1E). Furthermore, the use of a reporter virus reduces the turnaround time of virus detection by simply assessing luciferase values from five days to 24-48 hours. The NLuc signals, even at low MOIs, were robust and stable; however, this also led the signal to bleed in neighboring wells. This limitation was overcome by including appropriate mock-infected controls.

In addition, we have confirmed the potential to use the reporter virus in high throughput drug screens through proof-of-principle analysis of inhibition by remdesivir. The design of our reporter construct was based on similar NLuc SARS-CoV-2 viruses used for rapid screening of antiviral drugs effective against COVID-19 (7). We have assessed and optimized the use of our molecular clones for use in high throughput drug screens and confirmed results similar to ones done in parallel using a clinical virus isolate (VIDO-01) when treated with remdesivir in a dose-dependent manner. The resulting EC_50_of remdesivir for VIDO-01, and WTFL were comparable at 4.95, and 4.65 µM respectively. Similarly, the EC_50_for NLucFL using titrations was calculated as 6.55 µM, whereas using the NLuc assay it was 6.15 µM. In conclusion, we successfully optimized the antiviral reporter assay using the NLucFL virus. Moreover, this further emphasized that the molecular clones can be effectively used as efficient counterparts to clinical isolates for testing drugs.

Since working in biosafety level (BSL) 3 facilities to study SARS-CoV-2 is expensive, cumbersome, and sometimes not available, we aimed to design a replication assay system that can be used in conventional BSL2. We designed the SARS-CoV-2 sub-genomic replicon (SGR), based on previous SGR constructs used to study and discover inhibitors for HCV and SARS-CoV (12, 13). While the SGR showed evidence of replication based on an increase in luciferase expression over time which peaked at 24 hours, the replication assays were hindered by replicon-induced CPE leading to a rapid decline in luciferase counts. Similar luciferase expression patterns were observed in both Vero76 and Huh7.5 cells but in general, the results using the replicon system were inconsistent. The inconsistent luciferase numbers could be a result of varying viable RNA yields and quality obtained from *in vitro* transcription. Thus, although accessible as a BSL2 system, the transient assays were found to be inefficient and difficult to reproduce. Furthermore, we were unable to make stably expressing cell lines in Vero76 and Huh7.5 cells through selection with G418. In another report successful selection of a BHK21 cell line stably harboring autonomously replicating SARS-CoV-2 replicon RNA replicon cells required two attenuating mutations in NSP1 to reduce virus induced cellular toxicity (14). Other strategies to improve the ease of use of replicon systems include the use of a CMV promoter to remove the need for the *in vitro* transcription step. and generation of virus particles through trans-complementation that can be used for single round of replication assays (15).

In addition, we have generated novel human lung cell lines that can be used to evaluate SARS-CoV-2 replication, virus-host interactions, and screen drugs. Our goal was to generate cell lines that support SARS-CoV-2 and that are also readily killed by the infection so that they can be used in genetic screens that rely on live/dead selection. We also wished to identify other cell lines supporting SARS-CoV-2 replication with less CPE such that they might be amenable to the development of stable replicon cells. We tested several human lung cell lines since they are most likely to mimic virus-host interactions in the lung and possibly provide insight into mechanisms of lung pathogenesis. We also used HEK293T, a human kidney cell line which was permissive for SARS-CoV-2 replication as a positive control (16), and A549 a commonly used lung cell line. All the cell lines tested required transduction with ACE2 to become permissive to SARS-CoV-2. Three cell lines, NCI-H23^ACE2^, A549^ACE2^, and HEK293T^ACE2^, were notably successful in supporting virus infection while exhibiting more than >70% cell death post-infection (Figure 3). Clones from these cell lines were also generated and NCI-H23^ACE2^ clone A3 exhibited nearly 100% CPE following infection making it ideal for genetic screens such as CRISPR knockout and activation screens (Figure 3A-I). The HEK293T^ACE2^ monoclonal cell line clone A2, (Figure 3B-I) displayed CPE of approximately 90% following infection by SARS-CoV-2. These cell lines also showed robust NLuc expression after infection with SARS-CoV-2 NLucFL (Figure 3D). The monoclonal cell line A549^ACE2^ clone B1 showed about 70% CPE on day 4 post-infection (Figure 3C-I) with SARS-CoV-2 VIDO-01 isolate as well as increasing NLuc expression with SARS-CoV-2 NLucFL infection (Figure 3D). However, flow cytometry data suggest that these cells expressed less ACE2 (Figure 3C-II), and this may explain the delayed and less dramatic virus-induced cell death, but these cells could further be used for drug screening.

Transduction by ACE2 also allowed SARS-CoV-2 to infect NCI-H1703, NCI-H226, NCI-H520 and HCC827. Of these, NCI-H1703 ^ACE2^ exhibited ∼80% CPE. This cell line can be further looked at if another novel cell line is required to study SARS-CoV-2. Other cell lines transduced with ACE2 did not exhibit high CPE or overexpression of ACE2. Interestingly, these cell lines supported the growth of SARS-CoV-2 NLucFL assessed by NLuc expression as compared to their untransduced counterparts (Figure 4). These cell lines could be useful to study infections leading to less CPE.

Thus, we have generated viruses and tools that are valuable for the study of SARS-CoV-2 virus-host interactions and NCI-H23^ACE2^ clone A3 has already been used to assess the effects of an inhibitor on virus replication (9). These molecular clones can be further extended to study the impact of specific mutations on the phenotypes of SARS-CoV-2 variants of concern.

## Materials and Methods

### Cell lines and maintenance

All the cells were maintained at 37ºC with 5% CO_2_. HEK293T cells were cultured in Dulbecco’s modified Eagle medium (DMEM, with Sodium pyruvate) (HyClone #SH30243.01) supplemented with 10% fetal bovine serum (FBS) (Gibco #12483020) and 1× Penicillin-Streptomycin (PenStrep) (Gibco # 15140122). Vero76 cells were cultured in DMEM (without Sodium pyruvate) (Sigma #D5796) supplemented with 10% FBS and 1× PenStrep. A549 cells were a gift from Dr. Yan Zhou and were cultured in F-12K Medium (Kaighn’s Modification of Ham’s F-12 Medium) (ATCC #30-2004) supplemented with 10% FBS and 50μg/mL Gentamycin Sulfate (BioBasic #BS724). NCI-H23, NCI-H1703, HCC827, NCI-H226, and NCI-H520 were a gift from Dr. Deborah Anderson. The cells were cultured in Roswell Park Memorial Institute (RPMI) 1640 medium (Gibco #11875093) supplemented with 10% FBS and 1× PenStrep. The cryomedia used for freezing the cells contained 45% complete media, 45% FBS and 10% DMSO (MedChemExpress # HY-N7060). To test for mycoplasma contamination in all the cell lines, Mycoalert, Mycoplasma detection kit (Lonza #LT07-318) was used as per the manufacturer’s protocol.

### Virus stocks

Virus handling and related experiments were performed in the containment level 3 facility at Vaccine and Infectious Disease Organization (VIDO, SK, Canada). Most experiments with wild type virus were done using the P3 (passage #3) virus stock of SARS-CoV-2/Canada/ON/VIDO-01/2020 (hereafter referred to as SARS-CoV-2 VIDO-01) To prepare virus working stock, Vero76 cells were infected with the P2 virus at an MOI of ∼0.01. Briefly, virus stock was diluted in 10mL DMEM supplemented with 2% FBS and 1× PenStrep and inoculated in a T175 flask of 80% confluent Vero76 cells seeded 24 hours before infection. This was incubated at 37°C for 1 hour, gently rocking the flasks intermittently. After 1 hour, 25mL of complete media was added to the flask followed by incubation at 37ºC for 3-4 days until cytopathic effect (CPE) was observed. Once there was sufficient virus-induced CPE (∼70-80%), indicated by cell rounding and cell death, the supernatant was collected, centrifuged at 4000g for 10min, aliquoted in cryovials and stored at -80°C. The titer was determined by TCID_50_(as described below).

### Virus titration using TCID_50_

Vero76 cells were seeded 24 hours prior to infection in 96 well plates, such that they were 80% confluent the next day (1×10^4^ cells/ well). The virus to be titered was serially diluted 10-fold in a round bottom 96-well plate in DMEM supplemented with 2% FBS and 1 × PenStrep. 50µL volume of this was then inoculated in the seeded Vero76 cells and incubated at 37ºC for 1 hour. The virus inoculum was then removed and replaced with 100µL DMEM supplemented with 2% FBS and 1 × PenStrep. The plates were incubated at 37ºC, and CPE was noted by microscopy at 3-5 days post-infection. Titers were calculated using the Spearman-Karber algorithm(17).

### Virus titration using plaque assay

24 hours prior to infection, Vero76 cells were seeded in 12-well plates at 2.5 × 10^5^ cells/well. 400µL volume of 10-fold serially diluted virus (in DMEM supplemented with 2% FBS and 1 × PenStrep) was inoculated per well and incubated at 37ºC for 1 hour, with intermittent gentle rocking. After infection, the inoculum was replaced with 1.5mL 0.75% carboxymethyl cellulose (CMC) (Sigma #21902) media and incubated at 37ºC. To prepare the plaquing media, autoclaved CMC was dissolved in plain DMEM (Sigma #D5796) by placing it on a magnetic stirrer overnight at 4°C. Once the plaques of sufficient size developed (∼72 hours), the cells were fixed by adding 1.5mL 10% buffered formalin (Sigma #HT501128-4L) over the plaquing media and incubating at room temperature for 30min. The formalin-containing waste was discarded in Formalex ® Green formalin neutralizer (Jones Scientific #H-FORMG-CB). The plates were washed with water twice and then stained with 0.1% crystal violet [10% ethanol, 0.1% crystal violet, in water] for 30min. The stain was washed with water and the visible plaques were counted to calculate the titer.

### Design of SARS-CoV-2 molecular clones

Full-length SARS-CoV-2 Wuhan-1 (Wuhan seafood market pneumonia virus (2019-nCov) sequence NC_045512) cDNA genome cloned in a bacterial artificial chromosome (BAC) backbone was synthesized by CODEX DNA, Inc. The wild type full length (WTFL) (Codex Inc #SC2-FLSG-3333) construct is driven by the T7 promoter upstream of 5’ UTR, contains the D614G mutation and was designed to have a unique SbfI enzyme restriction site downstream of 3’ UTR and poly-A. For making the reporter virus expressing nanoluciferase (NLuc) (NLucFL) (Codex Inc; custom order), the NLuc gene was inserted by replacing the complete ORF7a while keeping the respective TRS sequence intact (7). For making the SARS-CoV-2 sub-genomic replicon (SGR) (Codex Inc; custom order), all the genes from spike (S) through ORF8 were replaced with Renilla luciferase (RLuc), Ubiquitin, and Neomycin resistance genes respectively, based on a SARS-CoV replicon (13). A plasmid encoding the Nucleocapsid gene (N gene) was a gift from Dr. Qiang Liu (18). Briefly, the codon-optimized N sequence is flanked by a T7 promoter on the 3’ end, and a 3xFLAG tag on the 5’ end, directly followed by a unique XbaI restriction site.

### *In vitro* RNA transcription

For making *in vitro* transcribed RNA, 2.5µg of WTFL and NLucFL or 1.5µg of SGR purified genomic DNA was linearized with 1.5µL of SbfI-HF^®^ (New England Biolabs #R3642L) in 1 × CutSmart buffer at 37ºC for 1 hour, followed by addition of 1.5 µL of SbfI-HF^®^ again for 1 hour. To remove the 3’ overhangs, the DNA was further treated with 1µL of Mung Bean Nuclease (New England Biolabs #M0250L) for 1 hour at 37ºC. The linearized DNA was extracted using phenol/chloroform and centrifugation in MaXtract High-density phase separation tubes (Qiagen #129046), followed by overnight precipitation with ammonium acetate and 100% ethanol at -20ºC. The next day the DNA was centrifuged at 14,000 rpm at 4°C for 30 minutes and the DNA pellet was dissolved in 6µL of nuclease-free water. This linearized DNA was *in vitro* transcribed to RNA as per the mMessage mMachine^™^ T7 kit protocol (Invitrogen #AM1344) with the following modifications. The total reaction volume was 50µL made up by adding the linearized DNA template with 7.5µL GTP (cap analog-to-GTP ratio of 1:1), 25µL 2X NTP/CAP, 5µL 10X reaction buffer, and 5µL enzyme mix. The reaction was incubated at 32ºC for 5 hours, followed by DNase treatment twice by adding 1µL Turbo DNase (provided in the kit) and incubating for 20 minutes at 37ºC each time. The RNA was extracted using phenol/chloroform and centrifuged in phase separation tubes and precipitated using isopropanol (7). To prepare the N mRNA, 1µg N gene plasmid DNA was linearized with XbaI enzyme for 1 hour at 37ºC, followed by Mung Bean Nuclease treatment for 1 hour at 37ºC. The purified and precipitated DNA was then used with mMessage mMachine^™^ T7 Ultra kit (Invitrogen #AM1345) according to the manufacturer’s protocol, this gave us *in vitro* transcribed N gene mRNA with an added poly-A tail to mimic the SARS-CoV-2 subgenomic mRNA. To quantify the *in vitro* transcribed (IVT) RNA for SARS-CoV-2 SGR and N gene, Qubit™ RNA High Sensitivity (HS) kit (Invitrogen # Q32852) was used as per the manufacturer’s protocol. We recovered ∼4-5μg of SARS-CoV-2 SGR RNA from 1.5μg of DNA template.

### Electroporation and virus production

To make viruses from the *in vitro* transcribed RNA, SARS-CoV-2 RNA and N gene RNA were electroporated in Vero76 cells. Briefly, cells were trypsinized and washed (with PBS) by centrifugation at 1000rpm for 5min. The cells we resuspended in Ingenio^®^ Electroporation Solution (Mirus Bio #MIR50114) such that the cell concentration was at 8 × 10^6^ per electroporation in 800μL final volume. 1-2 preps of *in vitro* transcribed RNA and 5-10µg N gene RNA were mixed with the prepared cell suspension in a 4mm electroporation cuvette (VWR International # 89047-210) and pulsed in a BioRad GenePusler using the exponential protocol with the following settings: Voltage-270V, Capacitance-950µF, Resistance - ∞ (infinity). Electroporated cells were equilibrated at room temperature for 5 minutes and transferred to a T-75 flask with 12 mL DMEM supplemented with 10% FBS and 1X PenStrep. (Alternatively, 1 prep of *in vitro* transcribed RNA with ∼5µg N gene RNA can be electroporated in 4 × 10^6^ cells in 400μL final volume and plated in a T-25 flask with 5mL DMEM supplemented with 10% FBS and 1 × PenStrep.) The electroporated cells were incubated at 37°C and monitored daily until significant CPE was observed 2-3 days post-electroporation. The virus stock was harvested as described above; this was considered as passage 0 (P0) stock. To passage the virus, 1mL of P0 virus stock was inoculated in a T-175 flask with 80% confluent Vero76 cells seeded 24 hours before infection. Once sufficient CPE was observed, the supernatant was harvested and stored as P1 stock.

For transient replication assays with SARS-CoV-2 SGR RNA, 4 million Vero76 cells resuspended in 400 μL of Ingenio^®^ Electroporation Solution were similarly electroporated with 1 prep of *in vitro* transcribed SGR RNA (prepared using 1.5 µg of SGR genomic DNA) and 5 µg of N gene RNA. After equilibrating at room temperature for 5 minutes, 100 μL of the electroporated cells were seeded in each well of a 6-well dish with 2 mL of DMEM supplemented with 10% FBS + 1X Pen-Strep. The cells were incubated at 37°C for 3 days and 100 μL media was harvested daily to assess Renilla luciferase expression. For harvesting cells from 6 well dishes, the growth medium was removed, and cells were rinsed with PBS. 200 μl/well of 1X Renilla lysis buffer was added to the well, and the cells were scraped off. The cell extracts were transferred to a microcentrifuge tube and stored at -80ºC until the luciferase assay was performed.

### Reporter luciferase assays

The Nano-Glo^®^ luciferase assay system (Promega #N1120) was used to assess NLuc expression by SARS-CoV-2 NLucFL, as per the manufacturer’s protocol. Briefly, virus infected 96-well plates were equilibrated at room temperature for ∼5-10 min. The Nano-Glo^®^ Luciferase Assay Reagent was prepared by combining one volume of Nano-Glo^®^ Luciferase Assay Substrate with 50 volumes of Nano-Glo^®^ Luciferase Assay Buffer and was also equilibrated to room temperature. 100 µL of the reagent was added to each of the wells already containing 100 µL infected media. The components were mixed well, incubated at room temperature for 3 minutes, and then transferred to white cell culture plates (Corning #C3610). Luminescence was measured using the Promega^™^ GloMax ® Explorer plate reader at 5 seconds of integration time. Luciferase assays to analyze SGR RNA replication was performed using the Renilla luciferase assay system (Promega #E2810) as per the manufacturer’s protocol with the following modifications. Before measuring the luminescence, cell extracts were centrifuged at 13000 rpm for 10 minutes at 4ºC, and 10 μL of the solution was mixed with 30 μL of Luciferase Assay Reagent. Luminescence was measured in a Promega^™^ GloMax^®^ 20/20 Luminometer (Promega #E5311) with an integration time of 10 seconds.

### Viral growth kinetics

To characterize the growth kinetics of patient and clone-derived SARS-CoV-2 stocks, supernatants from infected Vero76 cells were collected at different times post-infection and titered using TCID_50_. Briefly, cells were seeded 24 hours prior to infection in 96 well plates at 1 × 10^4^ cells/ well; for infection, the virus was diluted in DMEM supplemented with 2% FBS and 1 × PenStrep at MOI 0.01 (50 µL infection volume/ well). The cells were infected for 1 hour at 37ºC, after which the 50µL virus inoculum was replaced with 100µL fresh media (DMEM supplemented with 2% FBS and 1 × PenStrep). This marks the 0-hour post-infection timepoint. The supernatant was collected at 0, 2, 4, 6, 8, 24, 48, and 72 hours post-infection. For harvesting at different time points, 100 µL supernatant from respective wells was collected in a round bottom 96-well plate, sealed with a sealing tape (Nunc #12-565-398) and stored at -80ºC in Ziploc bags, until to be used for TCID_50_. The TCID_50_was performed in Vero76 cells, in 3-4 replicates. For assessing the luciferase kinetics, an identical 96-well plate was set up for infection in white cell culture plates (Corning #C3610). At the required time intervals, the Nano-Glo^®^ luciferase assay was performed as described above.

### Antiviral assay

To evaluate the antiviral effects of remdesivir on the wild type and reporter virus stocks, Vero76 cells were seeded 24 hours before infection in 96 well cell culture plates at 1 × 10^4^ cells/ well; for infection, the virus was diluted to a concentration of 1 × 10^2^ TCID_50_per 50µL in DMEM supplemented with 2% FBS and 1 × PenStrep. remdesivir (MedChemExpress #HY-104077) was reconstituted in DMSO to make the main stock. To generate a drug dose-response curve, remdesivir was 3-fold serially diluted in DMEM supplemented with 2% FBS, 1 × PenStrep and 0.1% DMSO. The cells were infected for 1 hour at 37ºC with 50µL of the diluted virus at an MOI 0.01, after which the virus inoculum was replaced with 100µL media containing the serially diluted remdesivir. The cells were incubated at 37ºC for 48 hours, and the viral supernatants were harvested and titrated using TCID_50_. The drug treatment was done in triplicates, followed by an equal number of TCID_50_for each of the replicates in Vero76 cells. In parallel, uninfected plates were treated with remdesivir to assess the cell viability under treatment. After 48 hours of drug treatment, the cell viability was assessed using the CellTiter 96^®^ AQ_ueous_One Solution Proliferation assay (Promega #G3580). Briefly, 20µL reagent was added to each well, incubated for 2 hours at 37ºC and read absorbance at 490nm as an endpoint assay on the Bio-Rad xMark^™^ Microplate Absorbance Spectrophotometer. For assessing the antiviral potency of compounds using the NLuc reporter system, an identical antiviral assay was set up in parallel, in white cell culture plates (Corning #C3610). After 48 hours of incubation, the Nano-Glo^®^ Luciferase assay was performed on the compound-treated infected wells as described above. The EC_50_of the compound and 50% cytotoxicity concentration (CC_50_) were determined in GraphPad Prism9 using the non-linear regression analysis. The cell viability was normalized to untreated cells depicting 100% viability, whereas the virus inhibition was normalized to untreated infected cells, depicting 100% virus luciferase expression.

### Generation of ACE-2 lentivirus particles

The lentivirus vectors containing the ACE2 gene were generated by co-transfecting psPAX2, pMD2.G, and EX-U1285-Lv197 (GeneCopoeia), an ACE2 gene containing lentivirus expression vector that also contained a Blasticidin selection gene. The plasmids were transfected into HEK293T cells using X-tremeGENE 9 (Roche # XTG9-RO) as per the manufacturer’s instructions. Eighteen hours post-transfection the media was replaced with DMEM containing 2% (w/v) bovine serum albumin (BSA) and then lentiviruses were collected after 24 and 48 hours (19).

### ACE2 transduction of cell lines and monoclonal cell selection

To make ACE2-expressing cell lines, ACE2 lentiviruses were filtered through a 0.45-micron filter and used to transduce respective cell lines using the reverse transduction method (Addgene, 2019). Briefly, 1-1.5 mL of filtered virus particles were added to the cell suspension of 50,000 cells in the appropriate media supplemented with 10% FBS and 8-10 μg/mL polybrene (Sigma # TR-1003-G) (without any antibiotics) and plated in 6 well cell culture plates with a final volume of 2.5-3 mL. 72 hours post-transduction, fresh media supplemented with 10% FBS and Blasticidin S HCl (Gibco # R21001) was added to cells. Cells were expanded in increasingly larger cell culture plates and ACE2 expression was confirmed by flow cytometry and based on susceptibility to infection with the SARS-CoV-2 VIDO-01 virus. The selection concentration of Blasticidin S HCl for HEK293T, and A549 cells was 5 μg/mL, for NCI-H23, HCC827 and NCI-H1703 was 4 μg/mL and for NCI-H520 and NCI-H226 was 2 μg/mL. After selection and 3-4 passages, the cells were maintained at half the selection concentration of Blasticidin S HCl. Single clone isolation from the HEK293T^ACE2^, A549^ACE2^ and NCI-H23^ACE2^ transduced cell pools was carried out by the array dilution method in 96-well plates (20). Cell colonies from single clones were collected 2-3 weeks after seeding and were expanded in increasingly larger cell culture plates.

### Analysis of cell surface ACE2 by flow cytometry

The cells to be analyzed were seeded in a 96-well plate at 1 × 10^5^ cells per well. At 70-80% confluency, healthy cells were detached from the monolayer using 0.5 mM EDTA (Gibco #15575020) in PBS and centrifuged at 1500 rpm for 3 min. The cell pellet was stained for 1 hour at 4°C with primary ACE2 antibody (R&D systems #AF933, used at a concentration of 0.25 µg/ 1 million cells). The cells were washed thrice with 100 µL flow wash buffer (2% FBS in PBS); followed by staining with secondary Goat IgG APC conjugated antibody (R&D systems # F0108, at the recommended volume of 10 µL/ 1 million cells) and 1000x live-dead viability stain (Invitrogen #L34958, used as 1 × final concentration) at 4ºC for 1 hour in the dark. The cells were washed with flow wash buffer before fixing with 2% paraformaldehyde (PFA) (diluted in flow wash buffer) for at least 20min or overnight. The cells were read on Beckman CytoFLEX Flow Cytometer.

### Infectivity assay to assess cell susceptibility

To assess whether the untransduced and ACE2 transduced cell lines were susceptible to SARS-CoV-2 infection leading to cytopathic effect, the cell lines were infected with SARS-CoV-2 VIDO and SARS-CoV-2-NLucFL. 24 hours before infection, 1-1.6 × 10^4^ cells were seeded in 96 well plates, such that they were 80% confluent the next day. The viruses were diluted to an MOI 0.5 in corresponding media supplemented with 2% FBS and 1x PenStrep, and the cells were infected for 1 hour at 37ºC with a volume of 50 µL virus inoculum. After infection, the virus inoculum was replaced with 100 µL fresh media supplemented with 2% FBS and 1x PenStrep. The CPE from SARS-CoV-2 VIDO-01 virus and NLuc expression from SARS-CoV-2 NLucFL was evaluated at 6, 24, 48 and 72 hours post-infection. The NLuc expression was assessed using the Nano-Glo® luciferase assay as described above. The virus-induced CPE was assessed using the Viral ToxGlo^™^ assay (Promega #G8943) according to the manufacturer’s protocol. Briefly, infected cells were equilibrated to room temperature for 5-10 min. One hundred microlitres of ATP detection reagent (prepared by adding ATP detection buffer to the ATP detection substrate) was added to the infected wells, incubated at room temperature for 10 min and the luminescence was read at 5 seconds integration time on a Promega^™^ GloMax ® Explorer plate reader. The luciferase values were normalized to blank readings (ATP detection reagent added to plain media).

### Data analysis and graphs

All data were analyzed and plotted in GraphPad Prism 9 software and the graphs are represented as mean +/- standard deviation.

## Acknowledgements

J.Q.K, and M.R, conceived the research and drafted the manuscript. J.Q.K, M.R, K.R, K.K.B, J.L, carried out the experiments. J.Q.K and M.R created the figures. All authors discussed the results and contributed to the revision of the final manuscript.

We thank Drs. Qiang Liu and Tom Hobman for providing plasmids, Dr. Deborah Anderson for providing an array of lung cell lines, Yan Zhou for A549 cells, Shelby Harms for assistance and training on SARS-CoV-2 antiviral assays, and Dr. Kerry Lavender, Madeline Stewart and Saurav Saswat Rout for their help with flow cytometry data analysis. This research was funded by a CIHR Operating Grant to JAW, FJV, and DF: COVID-19 Rapid Research Funding Opportunity – Therapeutics (VR3-172626). SARS-CoV-2 research is supported in the laboratory of D.F. by the Canadian Institutes of Health Research (CIHR; OV5-170349, VRI-173022 and VS1-175531). J.W. and D.F. are members of the CIHR-funded Coronavirus Variants Rapid Response Network (CoVaRR-Net). VIDO receives operational funding from the Government of Saskatchewan through Innovation Saskatchewan and the Ministry of Agriculture and from the Canada Foundation for Innovation through the Major Science Initiatives for its CL3 facility. JQK was funded by the College of Medicine (CoM) Devolved Scholarship and Graduate Research Fellowship (GRF) from the Biochemistry, Microbiology & Immunology Department, University of Saskatchewan. MR was funded by the College of Medicine (CoM) Devolved Scholarship and the Graduate Teaching Fellowship (GTF) from the Biochemistry, Microbiology & Immunology Department, University of Saskatchewan.

## References

1. Struyf T, Deeks JJ, Dinnes J, Takwoingi Y, Davenport C, Leeflang MM, Spijker R, Hooft L, Emperador D, Domen J. 2022. Signs and symptoms to determine if a patient presenting in primary care or hospital outpatient settings has COVID-19. Cochrane database of systematic reviews.

2. Hu Z, Song C, Xu C, Jin G, Chen Y, Xu X, Ma H, Chen W, Lin Y, Zheng Y. 2020. Clinical characteristics of 24 asymptomatic infections with COVID-19 screened among close contacts in Nanjing, China. Science China Life Sciences 63:706–711.

3. Syangtan G, Bista S, Dawadi P, Rayamajhee B, Shrestha LB, Tuladhar R, Joshi DR. 2020. Asymptomatic SARS-CoV-2 Carriers: A Systematic Review and Meta-Analysis. Front Public Health 8:587374.

4. V’Kovski P, Kratzel A, Steiner S, Stalder H, Thiel V. 2021. Coronavirus biology and replication: implications for SARS-CoV-2. Nat Rev Microbiol 19:155–170.

5. Naqvi AAT, Fatima K, Mohammad T, Fatima U, Singh IK, Singh A, Atif SM, Hariprasad G, Hasan GM, Hassan MI. 2020. Insights into SARS-CoV-2 genome, structure, evolution, pathogenesis and therapies: Structural genomics approach. Biochimica et Biophysica Acta (BBA)-Molecular Basis of Disease 1866:165878.

6. Zhang H, Penninger JM, Li Y, Zhong N, Slutsky AS. 2020. Angiotensin-converting enzyme 2 (ACE2) as a SARS-CoV-2 receptor: molecular mechanisms and potential therapeutic target. Intensive care medicine 46:586–590.

7. Xie X, Muruato A, Lokugamage KG, Narayanan K, Zhang X, Zou J, Liu J, Schindewolf C, Bopp NE, Aguilar PV, Plante KS, Weaver SC, Makino S, LeDuc JW, Menachery VD, Shi PY. 2020. An Infectious cDNA Clone of SARS-CoV-2. Cell Host Microbe 27:841–848 e3.

8. Park BK, Kim D, Park S, Maharjan S, Kim J, Choi J-K, Akauliya M, Lee Y, Kwon H-J. 2021. Differential signaling and virus production in Calu-3 cells and Vero cells upon SARS-CoV-2 infection. Biomolecules & therapeutics 29:273.

9. Ostrov DA, Bluhm AP, Li D, Khan JQ, Rohamare M, Rajamanickam K, k KB, Lew J, Falzarano D, Vizeacoumar FJ, Wilson JA, Mottinelli M, Kanumuri SRR, Sharma A, McCurdy CR, Norris MH. 2021. Highly Specific Sigma Receptor Ligands Exhibit Anti-Viral Properties in SARS-CoV-2 Infected Cells. Pathogens 10.

10. Hou YJ, Okuda K, Edwards CE, Martinez DR, Asakura T, Dinnon KH, 3rd, Kato T, Lee RE, Yount BL, Mascenik TM, Chen G, Olivier KN, Ghio A, Tse LV, Leist SR, Gralinski LE, Schafer A, Dang H, Gilmore R, Nakano S, Sun L, Fulcher ML, Livraghi-Butrico A, Nicely NI, Cameron M, Cameron C, Kelvin DJ, de Silva A, Margolis DM, Markmann A, Bartelt L, Zumwalt R, Martinez FJ, Salvatore SP, Borczuk A, Tata PR, Sontake V, Kimple A, Jaspers I, O’Neal WK, Randell SH, Boucher RC, Baric RS. 2020. SARS-CoV-2 Reverse Genetics Reveals a Variable Infection Gradient in the Respiratory Tract. Cell 182:429–446 e14.

11. Thi Nhu Thao T, Labroussaa F, Ebert N, V’kovski P, Stalder H, Portmann J, Kelly J, Steiner S, Holwerda M, Kratzel A. 2020. Rapid reconstruction of SARS-CoV-2 using a synthetic genomics platform. Nature 582:561–565.

12. Khan S, Soni S, Veerapu NS. 2020. HCV replicon systems: workhorses of drug discovery and resistance. Frontiers in Cellular and Infection Microbiology 10:325.

13. Hertzig T, Scandella E, Schelle B, Ziebuhr J, Siddell SG, Ludewig B, Thiel V. 2004. Rapid identification of coronavirus replicase inhibitors using a selectable replicon RNA. J Gen Virol 85:1717–1725.

14. Liu S, Chou CK, Wu WW, Luan B, Wang TT. 2022. Stable Cell Clones Harboring Self-Replicating SARS-CoV-2 RNAs for Drug Screen. J Virol 96:e0221621.

15. Malicoat J, Manivasagam S, Zuñiga S, Sola I, McCabe D, Rong L, Perlman S, Enjuanes L, Manicassamy B. 2022. Development of a Single-Cycle Infectious SARS-CoV-2 Virus Replicon Particle System for Use in Biosafety Level 2 Laboratories. Journal of virology 96:e01837–21.

16. Hoffmann M, Mösbauer K, Hofmann-Winkler H, Kaul A, Kleine-Weber H, Krüger N, Gassen NC, Müller MA, Drosten C, Pöhlmann S. 2020. Chloroquine does not inhibit infection of human lung cells with SARS-CoV-2. Nature 585:588–590.

17. Icho S, Rujas E, Muthuraman K, Tam J, Liang H, Landreth S, Liao M, Falzarano D, Julien JP, Melnyk RA. 2022. Dual Inhibition of Vacuolar-ATPase and TMPRSS2 Is Required for Complete Blockade of SARS-CoV-2 Entry into Cells. Antimicrob Agents Chemother 66:e0043922.

18. Nguyen HT, Falzarano D, Gerdts V, Liu Q. 2021. Construction of a noninfectious SARS-CoV-2 replicon for antiviral-drug testing and gene function studies. Journal of Virology 95:e00687–21.

19. Cunningham CE, Li S, Vizeacoumar FS, Bhanumathy KK, Lee JS, Parameswaran S, Furber L, Abuhussein O, Paul JM, McDonald M, Templeton SD, Shukla H, El Zawily AM, Boyd F, Alli N, Mousseau DD, Geyer R, Bonham K, Anderson DH, Yan J, Yu-Lee LY, Weaver BA, Uppalapati M, Ruppin E, Sablina A, Freywald A, Vizeacoumar FJ. 2016. Therapeutic relevance of the protein phosphatase 2A in cancer. Oncotarget 7:61544–61561.

20. Yan S. 2019. Genome editing: Isolating clones for genotypic and phenotypic characterization.

